# Data mining of Gene expression profiles of Saccharomyces cerevisiae in response to mild heat stress response

**DOI:** 10.1101/007468

**Authors:** Kaiyuan Ji, Wenli Ma, Wenling Zheng

## Abstract

We chose yeast as a model organism to explore how eukaryotic cells respond to heat stress. This study provides details on the way yeast responds to temperature changes and is therefore an empirical reference for basic cell research and industrial fermentation of yeast. We use the Qlucore Omics Explorer (QOE) bioinformatics software to analyze the gene expression profiles of the heat stress from Gene Expression Omnibus (GEO). Genes and their expression are listed in heat maps, and the gene function is analyzed against the biological processes and pathways. We can find that the expression of genes changed over time after heat stress. Gene expression changed rapidly from 0 min to 60 min after heat shock, and gene expression stabilized between 60 min to 360 min. The yeast cells begin to adjust themselves to the high temperatures in terms of the level of gene expression at about 60 min. In all of the involved pathways and biological processes, those related to ribosome and nucleic acid metabolism declined in about 15–30 min and those related to starch and sucrose increased in the same time frame. Temperature can be a simple way to control the biological processes and pathways of cell.

## 1. Introduction

The yeast monoplast is a model eukaryote for studies in molecular and cellular biology. It has been used in fundamental and applied research for many years and is a genetically tractable organism, amenable to modifications such as gene marking, gene disruption, gene mutation and gene-dosage effects. A comparison of the human and yeast genomes revealed that almost 30% of known genes involved in human disease have a homology within yeast [1]. Up to now, advances in genome-wide screening have facilitated the use of this model for medical research [2]. Cells grow optimally within a relatively small range of external stimuli but tolerate moderate deviations, some of which influence cell structure and function via rapid physiological adaptations. One of the adaptation mechanisms is the heat shock response, a highly conserved program of changes in gene expression that result in changes in a broad range of cellular processes, including transient arrest of cell division [3], uncoupling of oxidative phosphorylation [4], accumulation of misfolded proteins [5], and changes in the cell wall and membrane dynamics [6, 7]. Also heat shock restrains cell growth and viability, as well as anabolic and catabolic activity [8]. Yeast exhibits optimal growth between 25 °C to 30 °C. But at temperatures beyond 36–37°C, yeast activates a protective transcriptional program and alters other components of its physiology. In previous decades, numerous gene-specific investigations have been performed on the diverse pathways that are affected, and our knowledge of heat shock response is rich and detailed. We now have in hand a program of how expression changes during a time period. Since yeast RNA polymerase II is inactive at temperatures beyond 42 °C [9], we used the GSE25503 data from the Gene Expression Omnibus (GEO) to begin our data mining study, which focuses on the transcriptional response of S. cerevisiae to an increase in temperature from 28 °C to 41 °C. We analyzed GSE25503 conbine with GSE406 and GDS112 to insure the result more accurate and significative. These results will benefit both biological basic research and industrial fermentation.

## 2. Materials and methods

### 2.1 GEO data

Gene expression profile data GSE25503, GSE406, GDS112 were downloaded from the open Gene Expression Omnibus database (http://www.ncbi.nlm.nih.gov/geo/).

The GSE25503 data contains the whole-genome transcriptional response of S. cerevisiae to an increase in temperature from 28 °C to 41 °C under well-controlled conditions. Samples were taken from three replicate control cultures at 28°C at three points in time and from three replicate stressed cultures at six points in time. This data was uploaded by Femke Mensonides, University of Amsterdam.

The GSE406 data contains the whole-genome transcriptional response of wild type and kin82 mutant of S. cerevisiae to an increase in temperature from 25 °C to 37 °C. Samples were collected at 0, 5, 15, 30 and 60 minutes after transfer to 37°C. This data was uploaded by Eran Segal, Stanford University.

The GDS112 data is the whole-genome transcriptional response of S. cerevisiae which cultures shifted from 30°C to 37°C and samples collected at 0, 5, 15, 30 and 60 minutes. Data was uploaded by Stanford Microarray Database.

### 2.2 QOE analysis

The Qlucore Omics Explorer (QOE) software provides new technology for data analysis and data mining. It offers state-of-the-art mathematical and statistical methods, and its main features are the ease of use and speed with which data sets can be analyzed and explored. The analysis is supported using general statistical methods and measures, such as t-tests and F-tests with the corresponding p-values and false discovery rates. The GEO data can be immediately imported into QOE 3.0, and the respective annotations can also be downloaded.

When data is imported into QOE, we normalize it to a mean = 0 and variance = 1 for each variable. In order to uncover hidden structures and to find patterns in large data sets, our statistic analysis used dynamic Principal Component Analysis (dynamic PCA) and hierarchical clustering. We computed all the variance of variables (σ) and filter the variance by the ratio of σ/σ_max_. σ_max_ is the maximum of variance in all variables. Multi-group comparisons and two-group comparisons can help us find the available variables, and all data sets were identified according to the gene symbol and collapsed on average. The results are tested with the appropriate p-value and q-value. Because there are multiple repeat in each group of GSE25503.The variables of GSE25503 are analyzed by multi-group comparisons. Given there is one sample in each group, the variables of GSE406 and GDS112 are filtered by ratio of σ/σ_max_. We choose samples which collected in 0-60mins for uniformity and wild type only was used in our analysis. The intersection of gene list was obtained from these three data sets.

### 2.3 Biological process and pathway analysis

The Database for Annotation, Visualization and Integrated Discovery (DAVID) is a software package that uses a built-in graphical interface that gives access to an ample database. We used DAVID to identify KEGG categories with a p-value < = 0.1 and count >= 2 based on Fisher Exact statistics. The Saccharomyces Genome Database (SGD) provides comprehensive, integrated biological information for the budding yeast Saccharomyces cerevisiae along with search and analysis tools that can be used to explore the data, enabling the analysis of Gene Ontology (GO) biological process with a p-value < 0.01 [10].

## 3 Results

### 3.1 Multi group comparison of differentially expressed genes in 360mins

Dynamic PCA analysis reduces the dimensionality of the data in order to generate three-dimensional graphical representations. In this process, as much variance as possible from the original data is retained. In order to explore all of the gene expression changes in different time periods, groups at 120 min and 240 min, which had temperatures of 28 °C were sampled, and other groups were sampled to conduct multi-group comparison at different points in time. After performing the multi group comparison (p = 3.6569 e-8, q = 1.5 e-7), we identified 1390 variables. Dynamic PCA showed the structure of GSE25503 (Fig. 1), and we can find that groups at 60 min, 120 min, 240 min and 360 min were closer than others, which means they were similar in gene expression. Groups at 0 min, 10 min, and 30 min were quite difference, especially between 0 min and 30 min. The samples also showed a tendency for changes in gene expression over time – gene expression changed the most in the first 30 mins, and then it returned toward the status of approximately the 0-min group. To explore the detail in the first 60mins, we analyzed the same differentially expressed genes of GSE25503, GSE406 and GDS112.

**Figure 1:**
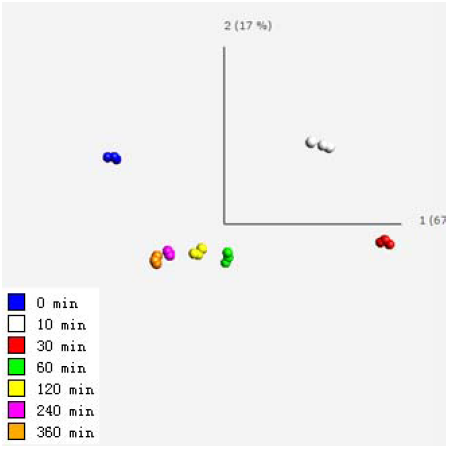
Dynamic PCA map of the GSE25503 data. The basic interpretation of the PCA plot of the data sets in QOE is that the data points that are similar with respect to the observed data values are also presented close together in the generated plots.

### 3.2 Same differentially expressed genes of GSE25503, GSE406 and GDS112 in 60mins

The 1000 most differentially expressed genes of GSE25503 ((p = 2.5304 e-6, q = 1.444 e-5), GSE406 (σ/σ_max_ = 0.23582) and GDS112 (σ/σ_max_ = 0.23125) in time were intersected. So we obtain 361 same differentially expressed genes. Dynamic PCA maps of 361 same differentially expressed genes of GSE25503, GSE406 and GDS112 are shown in Fig 2. We can find that there is a significant difference among each group. The samples form a circular pattern in three-dimensional space. The heat maps of the 361 differentially expressed genes are shown in Fig 3 after hierarchical clustering. The genes can falls into two parts due to hierarchical clustering. The genes of first part express low after heat stress and the genes of another part express high after heat stress. Most genes changed between 0 min and 60 min. At 60 min, the expression of some genes was similar to that of the 0-min group. The gene symbols of these genes are shown to the right of heat map, and the samples varied with respect to time.

**Figure 2:**
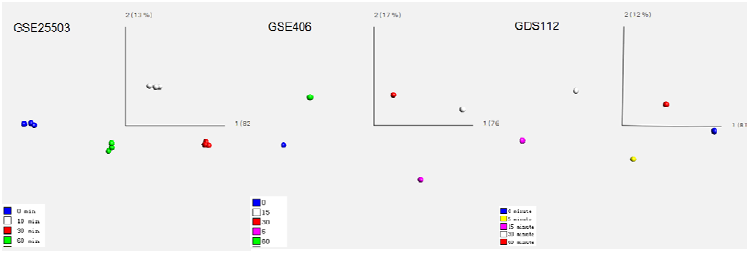
Dynamic PCA map of 361 same differentially expressed genes of GSE25503, GSE406 and GDS112 in 60mins.

**Figure 3:**
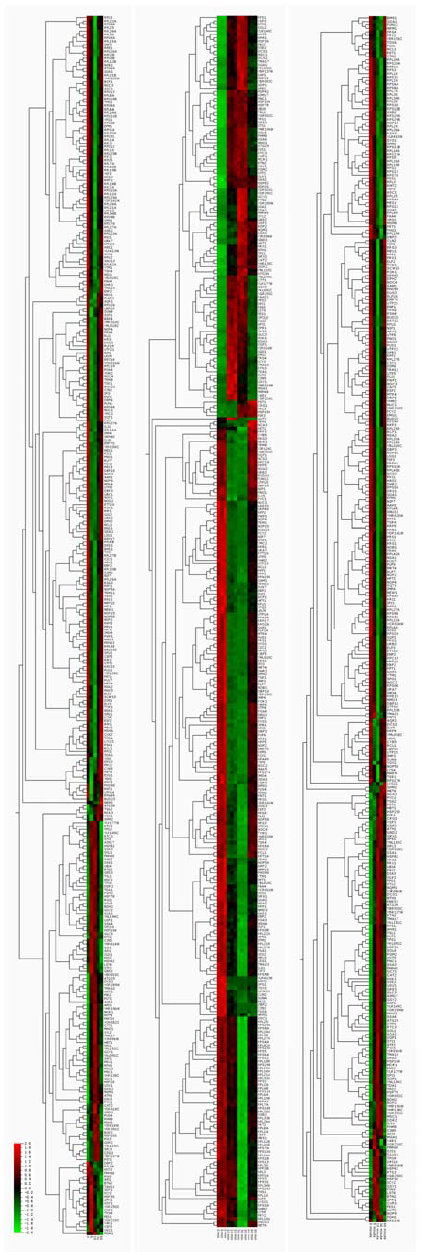
Heat map of 361 same differentially expressed genes. The red color indicates that the variable has a high value for that sample and green indicates that the variable has a low value for that sample.

The biological processes of 361 same differentially expressed genes are analyzed by SGD. The relationship of these biological processes is shown in Fig 4. GO term, GOID and P-value are listed in Table 1. Most of the involved biological processes are related to glycometabolism, ribosome and nucleic acid metabolism. The most heavily affected biological processe is ribosome biogenesis.

**Figure 4:**
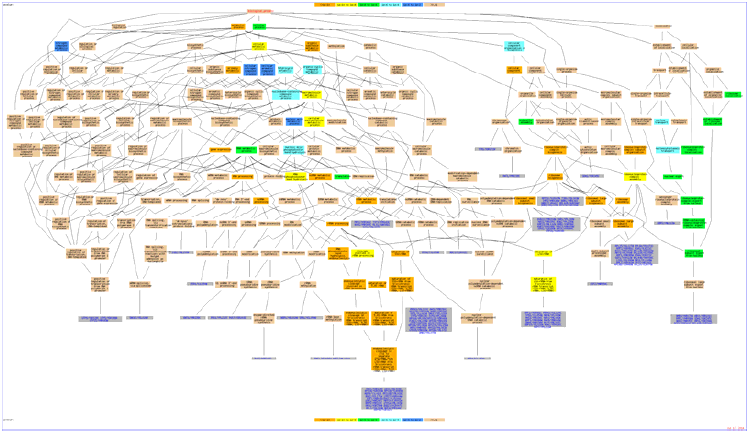
The map shows the biological processes of 361 same differentially expressed genes. Genes in the graphic are associated with the GO term(s) which are directly annotated. The color of each box indicates the p-value score. Genes associated with the GO terms are shown in gray boxes.

**Table 1:** The list of the biological process involved

| GOID | GO_term | P-value |
| --- | --- | --- |
| 42254 | ribosome biogenesis | 5.93E-62 |
| 22613 | ribonucleoprotein complex biogenesis | 5.20E-54 |
| 6364 | rRNA processing | 1.14E-37 |
| 16072 | rRNA metabolic process | 1.31E-36 |
| 34470 | ncRNA processing | 4.61E-36 |
| 34660 | ncRNA metabolic process | 9.18E-35 |
| 2181 | cytoplasmic translation | 2.39E-31 |
| 42273 | ribosomal large subunit biogenesis | 3.56E-29 |
| 42274 | ribosomal small subunit biogenesis | 2.54E-28 |
| 30490 | maturation of SSU-rRNA | 2.29E-25 |
| 462 | maturation of SSU-rRNA from tricistronic rRNA transcript (SSU-rRNA, 5.8S rRNA, LSU-rRNA) | 1.01E-23 |
| 6396 | RNA processing | 7.99E-23 |
| 44085 | cellular component biogenesis | 7.66E-21 |
| 42255 | ribosome assembly | 2.90E-16 |
| 460 | maturation of 5.8S rRNA | 5.57E-16 |
| 466 | maturation of 5.8S rRNA from tricistronic rRNA transcript (SSU-rRNA, 5.8S rRNA, LSU-rRNA) | 5.57E-16 |
| 44238 | primary metabolic process | 6.04E-15 |
| 71704 | organic substance metabolic process | 2.86E-14 |
| 8152 | metabolic process | 1.01E-13 |
| 10467 | gene expression | 1.56E-12 |
| 27 | ribosomal large subunit assembly | 4.57E-12 |
| 71826 | ribonucleoprotein complex subunit organization | 3.04E-11 |
| 478 | endonucleolytic cleavage involved in rRNA processing | 3.74E-11 |
| 479 | endonucleolytic cleavage of tricistronic rRNA transcript (SSU-rRNA, 5.8S rRNA, LSU-rRNA) | 3.74E-11 |
| 90502 | RNA phosphodiester bond hydrolysis, endonucleolytic | 5.95E-11 |
| 447 | endonucleolytic cleavage in ITS1 to separate SSU-rRNA from 5.8S rRNA and LSU-rRNA from tricistronic rRNA transcript (SSU-rRNA, 5.8S rRNA, LSU-rRNA) | 7.75E-11 |
| 44237 | cellular metabolic process | 1.74E-10 |
| 22618 | ribonucleoprotein complex assembly | 2.09E-10 |
| 43170 | macromolecule metabolic process | 2.75E-10 |
| 470 | maturation of LSU-rRNA | 5.00E-10 |
| 44260 | cellular macromolecule metabolic process | 5.12E-10 |
| 469 | cleavage involved in rRNA processing | 1.47E-09 |
| 90501 | RNA phosphodiester bond hydrolysis | 2.02E-09 |
| 463 | maturation of LSU-rRNA from tricistronic rRNA transcript (SSU-rRNA, 5.8S rRNA, LSU-rRNA) | 5.08E-09 |
| 9987 | cellular process | 1.06E-08 |
| 16070 | RNA metabolic process | 5.55E-08 |
| 70925 | organelle assembly | 7.56E-08 |
| 33750 | ribosome localization | 2.35E-07 |
| 33753 | establishment of ribosome localization | 2.35E-07 |
| 54 | ribosomal subunit export from nucleus | 2.35E-07 |
| 71166 | ribonucleoprotein complex localization | 2.35E-07 |
| 71426 | ribonucleoprotein complex export from nucleus | 2.35E-07 |
| 71428 | rRNA-containing ribonucleoprotein complex export from nucleus | 2.35E-07 |
| 51168 | nuclear export | 3.52E-07 |
| 6412 | translation | 6.13E-07 |
| 5991 | trehalose metabolic process | 4.26E-06 |
| 6913 | nucleocytoplasmic transport | 4.32E-06 |
| 34471 | ncRNA 5'-end processing | 4.48E-06 |
| 967 | rRNA 5'-end processing | 4.48E-06 |
| 51169 | nuclear transport | 4.96E-06 |
| 966 | RNA 5'-end processing | 6.69E-06 |
| 90305 | nucleic acid phosphodiester bond hydrolysis | 6.91E-06 |
| 6139 | nucleobase-containing compound metabolic process | 8.32E-06 |
| 1901360 | organic cyclic compound metabolic process | 1.83E-05 |
| 472 | endonucleolytic cleavage to generate mature 5'-end of SSU-rRNA from (SSU-rRNA, 5.8S rRNA, LSU-rRNA) | 3.42E-05 |
| 46483 | heterocycle metabolic process | 6.49E-05 |
| 71840 | cellular component organization or biogenesis | 9.55E-05 |
| 6725 | cellular aromatic compound metabolic process | 0.00011 |
| 480 | endonucleolytic cleavage in 5'-ETS of tricistronic rRNA transcript (SSU-rRNA, 5.8S rRNA, LSU-rRNA) | 0.00012 |
| 90304 | nucleic acid metabolic process | 0.00023 |
| 19321 | pentose metabolic process | 0.00079 |
| 51029 | rRNA transport | 0.00083 |
| 6407 | rRNA export from nucleus | 0.00083 |
| 34641 | cellular nitrogen compound metabolic process | 0.00095 |
| 6807 | nitrogen compound metabolic process | 0.00157 |
| 46351 | disaccharide biosynthetic process | 0.00468 |
| 5992 | trehalose biosynthetic process | 0.00468 |
| 9312 | oligosaccharide biosynthetic process | 0.00468 |

We used DAVID to identify KEGG PATHWAY categories with a modified Fisher Exact P-Value < 0.1. There are 4 pathways involved, and the ribosomal pathway’s p-value was the lowest of all (as shown in Table 1). All of the heat maps of the genes which are in the pathway list below are displayed in Fig 4. The gene symbols of these genes are shown to the right of heat map, and the samples varied with respect to time. Most of ribosome pathway genes express low in 15-30 min and some of them begin to express low since 5-10 min, such as RPL24, RPL7 and RPS27A. In starch and sucrose metabolism pathway, almost all the genes express low since 5-10 min except EXG1. PRS1, PRS3, PRS4 and RKI1 express low in 5-30 min after heat stress, and others express oppositely. All the pentose and glucuronate interconversions involved genes express high in 5-30 min.

**Figure 5:**
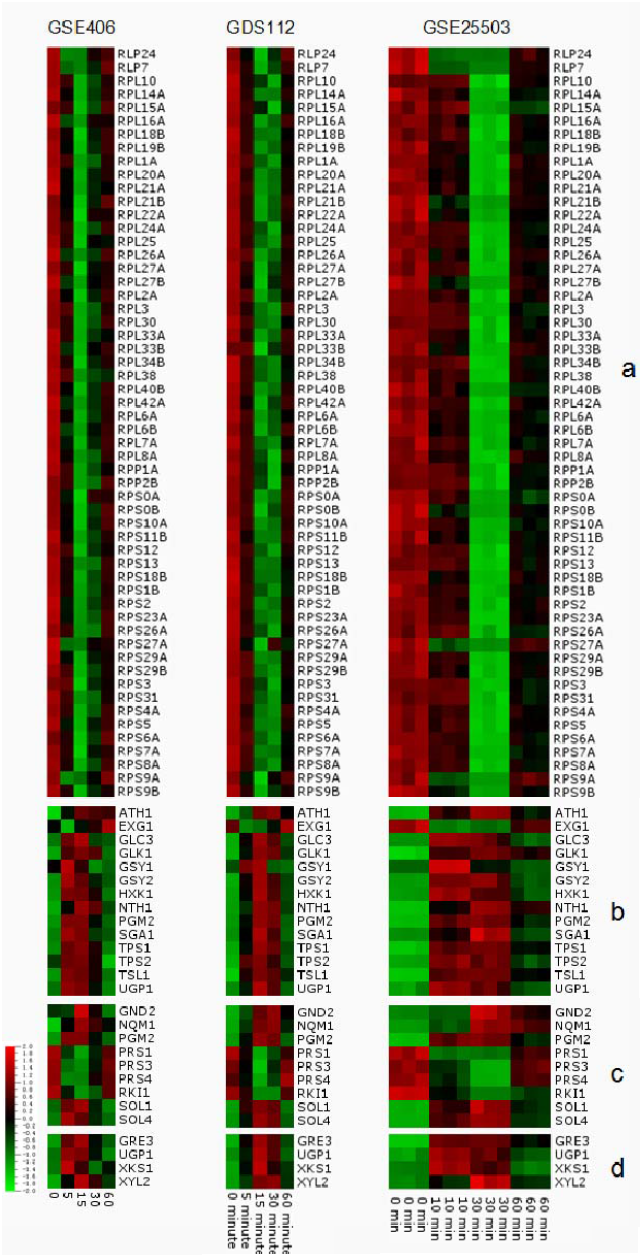
Heat map of 361 same differentially expressed genes. a: Ribosome pathway; b: Starch and sucrose metabolism pathway; c: Pentose phosphate pathway; d: Pentose and glucuronate interconversions.

**Table 2:**
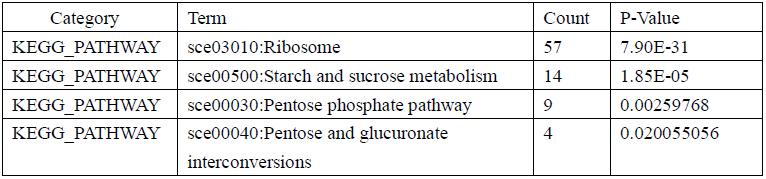
Enriched pathways of 1390 differentially expressed genes.

| Category | Term | Count | P-Value |
| --- | --- | --- | --- |
| KEGG_PATHWAY | sce03010:Ribosome | 57 | 7.90E-31 |
| KEGG_PATHWAY | sce00500:Starch and sucrose metabolism | 14 | 1.85E-05 |
| KEGG_PATHWAY | sce00030:Pentose phosphate pathway | 9 | 0.00259768 |
| KEGG_PATHWAY | sce00040:Pentose and glucuronate interconversions | 4 | 0.020055056 |

## 4 Discussion

### 4.1 **Y**east cells begin to adjust the level of gene expression to the high temperature at about 60 min.

After heat shock, we can see that all the samples form a ring-like 3D-PCA map (Fig 1). The samples at 60 min, 120 min, 240 min and 360 min are close to each other. However, the distances of the samples from 0 min to 10 min, 10 min to 30 min and 30 min to 60 min are long. As we know, the distance between the samples signifies a difference in gene expression between them. Therefore, we can learn from the PCA map that the level of gene expression changes rapidly from 0 min to 60 min after heat shock, and the level of gene expression changed slightly from 60 min to 360 min. There were considerable genetic differences between the 0-min group and the 30-min group. After 30 min, the variation in the trend returned to a state approaching that of the 0-min group, but some distance remained. The stability of the gene expression after 60 min indicates that yeast cells begin to adjust to the high temperature with respect to the level of gene expression.

### 4.2 Biological Process and PATHWAY analysis of same differentially expressed genes

When the temperature is beyond that of a normal state, the cells can sense heat shock as a result of the accumulation of thermally misfolded proteins [11]. Two major transcriptional control and regulation systems occur upon heat shock: those involving heat shock factors (HSF) and those involving MSN2 and MSN4 [12, 13]. These bind to the heat shock elements (HSR) and stress response element (STRE) [14, 15]. Yeast treated with a mild heat shock accumulates stored trehalose, which requires MSN2 and MSN4. MSN2/4 regulate the expression of hundreds of genes in response to several stresses, including heat shock, osmotic shock, oxidative stress, low pH, glucose starvation, and high concentrations of sorbic acid and ethanol [16, 17]. HSF1 regulates protein folding, detoxification, energy generation, carbohydrate metabolism, and cell wall organization [18]. However, of all of the involved pathways, we can find that the ribosome biogenesis has the highest degree of enrichment. The heat map of the ribosome pathway shows that most ribosome proteins have the lowest expression at 30 min. In the early research, Yan Liu found that heat shock disassembles the nucleolus [19], which is the factory of rRNA [20]. Ribosomes are specialized complexes composed of of nucleic acids and proteins that are responsible for mediating protein synthesis. Nucleic acids, (rRNA and tRNA) molecules are essential for a ribosome to translate mRNA into proteins [21-23]. In these three data sets, the ribosome pathways showed a decline in the 15–30 min range.

The analysis of the biological processes shows that most of the enrichment processes is related to the metabolism of nucleic acids and ribosome. This must affect the growth and breeding of the yeast. In Pentose phosphate pathway and Yeast cells complete a cell cycle in approximately 70 to 90 min, and the key regulatory checkpoint starts in the G1-to-S-phase transition [24]. Heat shock leads to a transient arrest at precisely this stage in the cell cycle [25] due to the reduction of the transcript levels of the G1/S cyclins [26]. We thought this was also a result of the change of the pathways related to the nucleic acids to the amino acids.

Trehalose metabolic process is also an involved biological process. Trehalose is an important carbohydrate store in yeast, and the ability of cells to withstand heat shock correlates with accumulates of cellular trehalose [27]. The starch and sucrose metabolism pathway shows high levels of gene expression. At 10 min to 30 min, in we can find TPS1 and TPS2 genes that are encoded trehalose-6-phosphate synthase (TPS) and trehalose-6-phosphate phosphatase (TPP), and these are expressed higher in the starch and sucrose metabolism pathways [28]. The low expression of EXG1and high expression of other genes in starch and sucrose metabolism pathways would accelerate accumulate of trehalose and D-glucan which is the main construction of cell wall [29]. Combine Pentose and glucuronate interconversions pathway with Pentose phosphate pathway, we can find heat stress promote D-ribose produce in 5-30 min.

Heat shock can lead to thousands of changes in gene expression. Under mild heat stress, yeast can survive by adjusting gene expression. By identifying differentially expressed genes and the biological processes and pathways thereof, genes closely related to heat stress were screened. After about 60 min, gene expression of yeast tends to have stability. In other words, yeast adapts the high temperatures. However, environment and stimulate can affect the cell’s biological processes and pathways which will be our tools to control the cell.

